# How well can we create phased, diploid, human genomes?: An assessment of FALCON-Unzip phasing using a human trio

**DOI:** 10.1101/262196

**Authors:** Arkarachai Fungtammasan, Brett Hannigan

## Abstract

Long read sequencing technology has allowed researchers to create de novo assemblies with impressive continuity[1,2]. This advancement has dramatically increased the number of reference genomes available and hints at the possibility of a future where personal genomes are assembled rather than resequenced. In 2016 Pacific Biosciences released the FALCON-Unzip framework, which can provide long, phased haplotype contigs from de novo assemblies. This phased genome algorithm enhances the accuracy of highly heterozygous organisms and allows researchers to explore questions that require haplotype information such as allele-specific expression and regulation. However, validation of this technique has been limited to small genomes or inbred individuals[3].

As a roadmap to personal genome assembly and phasing, we assess the phasing accuracy of FALCON-Unzip in humans using publicly available data for the Ashkenazi trio from the Genome in a Bottle Consortium[4]. To assess the accuracy of the Unzip algorithm, we assembled the genome of the son using FALCON and FALCON Unzip, genotyped publicly available short read data for the mother and the father, and observed the inheritance pattern of the parental SNPs along the phased genome of the son. We found that 72.8% of haplotype contigs share SNPs with only one parent suggesting that these contigs are correctly phased. Most mis-phased SNPs are random but present in high frequency toward the end of haplotype contigs. Approximately 20.7% of mis-phased haplotype contigs contain clusters of mis-phased SNPs, suggesting that haplotypes were mis-joined by FALCON-Unzip. Mis-joined boundaries in those contigs are located in areas of low SNP density. This research demonstrates that the FALCON-Unzip algorithm can be used to create long and accurate haplotypes for humans and identifies problematic regions that could benefit in future improvement.

## INTRODUCTION

Long genomic reads generated using Pacific Bioscience’s single-molecule, real-time sequencing technology can be used to generate reads with high consensus accuracy[1,2]. This technology has significantly enhanced our ability to create de novo assemblies with high contig continuity. Long reads are capable of bridging over repetitive regions[5,6,2,7] which are particularly problematic for reads from short-read technologies[8,5,9–11,7,12], and they can be used to create assemblies at much higher throughput and lower cost than traditional Sanger sequencing[6,13]. These long reads also do not have coverage bias due to skewed G+C content [7,14], a phenomenon particularly prevalent in microbial genomes[12,15,16]. These advancements have dramatically increased the number of high-quality reference genomes across multiple taxa, greatly benefiting the fields of comparative genomics and metagenomics[2,7,12].

To enhances the accuracy of *de novo* genome assemblies of highly heterozygous diploid organisms, Pacific Biosciences released the FALCON-Unzip[3] framework. After de novo genome assembly by FALCON, the Unzip framework can be used to create long, phased haplotype contigs (haplotigs) which represent the variants found along a single copy of a pair of homologous chromosomes in a diploid organism. This phasing information can be used to study allele-specific expression, identify compound heterozygous mutations, or study population differentiation [17,18,7]. Phased haplotypes also offer insight into low-frequency and de novo variants which may not be reliably inferred through statistical methods that use population haplotypes unless individuals related to the individual are also genotyped[17]. If de novo assembly can provide an accurately phased genome, these benefits could argue for selecting a de novo assembly approach rather than a resequencing approach for studying human genetic variation.

However, to date, evaluations of the phasing accuracy of FALCON-Unzip have used smaller genomes, and inbred species such as Arabidopsis[3]. It is of great interest to the scientific community to also evaluate FALCON Unzip’s accuracy on larger genomes, genomes with more repetitive elements[19–21], or genomes with complex structure.

As a roadmap to personal genome assembly and phasing, we chose to evaluate the accuracy of FALCON-Unzip phasing in humans. We performed de novo genome assembly and phasing for the son in the Ashkenazi trio from the Personal Genomes Project [4] using the FALCON and FALCON Unzip pipelines. Using high coverage Illumina data, we identified SNPs that can be traced back to the mother or father which could be used to identify the parental origin of a given haplotig. We then asked two questions: first, what percentage of haplotigs contain SNPs entirely consistent with a single parent, and second, what patterns are observed in haplotigs with SNPs from both parents.? These patterns may provide insight to researchers improving upon the underlying phasing algorithms.

## MATERIALS AND METHODS

### Implementation of Falcon and Falcon-Unzip in the cloud environment

The genome assemblies were performed on the DNAnexus cloud platform using the FALCON 1.7.7 pipeline from Pacific Bioscience[22], FALCON Unzip from the master branch of the Unzip GitHub repo retrieved in January 2017, and the Damasker suite from Gene Myers[23]. These apps are available for public use by request.

### Falcon genome assembly and Falcon-Unzip phasing of Ashkenazi

First, 70-fold whole-genome, single-molecule, real-time sequencing (SMRT) data of a male Ashkenazi (HG002) was provided by the Genome in a Bottle Consortium hosted by the National Institute of Standards and Technology[4]. This data was passed through the TANmask and REPmask modules from the Damasker suite. These tools mask repetitive regions from use during the initial read overlap discovery, causing the overlapper to instead focus on unique sequences to identify reads that overlap because they come from the same genomic locus. This masked data was then used as input to the traditional FALCON pipeline, using a length cut-off of 8,304 bp which represents the longest contigs that provide 50x coverage of the human genome during the initial overlap stage which will be used to error-correct the original raw reads.

Second, the error-corrected reads were then passed through the TANmask and REPmask modules, followed by the overlap portion of the FALCON pipeline. For this second overlap portion, a length cut-off of 9,778 bp was selected, representing the longest contigs that provide 20x coverage of the human genome.

Finally, the assembled genome was separated into haplotype contigs (haplotigs) using FALCON-Unzip[24], which includes a polishing step that leverages PacBio’s Quiver algorithm from SMRT Link 4[25]. The outputs from FALCON-Unzip consist of primary contigs and haplotigs. The primary contigs are continuous sequences that represent the assembled haploid genome. For the substring of a primary contig that has sufficient evidence of variation between two homologous chromosomes, there will be a haplotig which represents the haplotype of one parent while the primary contig will contain the haplotype of the other parent (Fig 1A). Each primary contig could be composed of a mixture of segments from both parents, but the haplotig and the segment of a primary contig that corresponds to that haplotig should contain only variants from a single parent if the phasing algorithm performs correctly.

**1.**
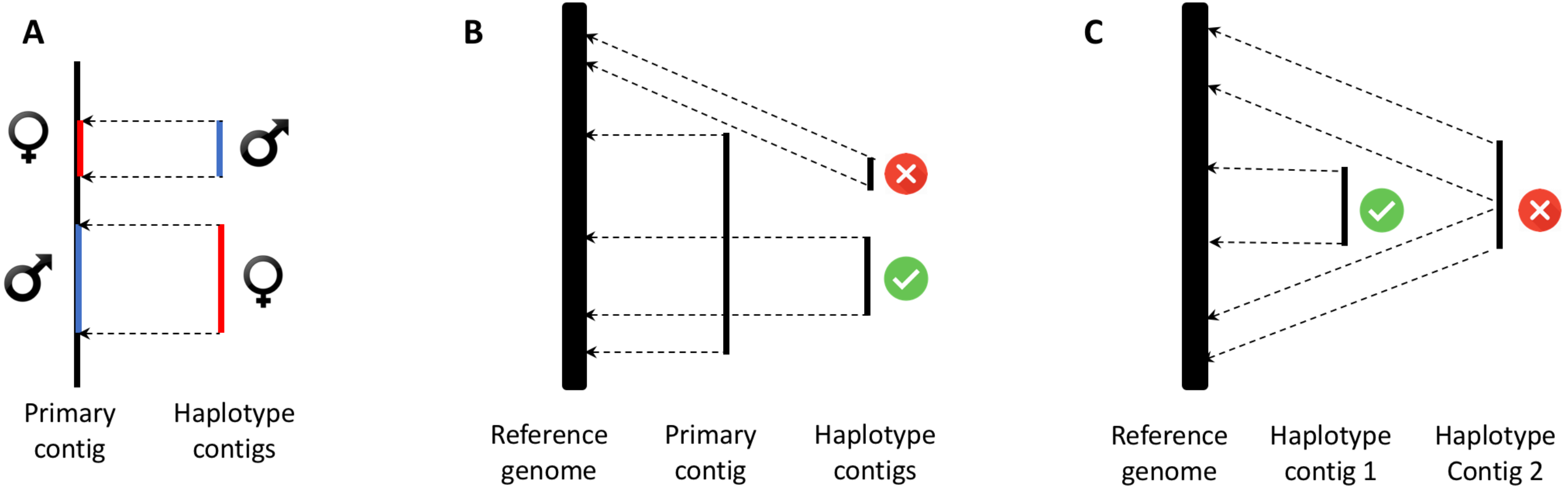
Screening of primary and haplotype contigs. A) Primary and haplotype contigs. B) Removing a haplotype contig that mapped outside its corresponding primary contig. C) Removing a haplotype contig that were interrupted by another haplotig along reference genome mapping.

### Coordinating and calling variants of primary and haplotype contigs

In order to impose a coordinate system on our new assembly, the primary contigs were mapped to the GRCh38 human reference genome[13] using the nucmer tool from the MUMmer3.23 package[26]. Mappings were filtered using MUMmer’s delta-filter screening tool to remove any part of primary contig that mapped to more than one location in the reference genome. We then used MUMmer’s show-snps function to call variants on the uniquely mapped primary contigs. Using the same mapping, filtering, and variant calling procedure, we produced a set of variant calls for the haplotigs as well.

We observed that in some instances a haplotig would fail to map to the same genomic region as its corresponding primary contig (Fig 1B). This discordance could be due to a few issues. First, haplotigs are shorter than primary contigs since by definition they are substrings of the primary contigs for which phasing can be determined. Due to the decreased length of haplotigs, they are more prone to mapping errors, especially at repetitive regions or sites of structural rearrangement. On the other hand, primary contigs are longer and may contain assembly errors such as incorrectly combining non-adjacent segments of DNA into the same contig or placing two neighboring pieces of DNA in the incorrect order. These assembly errors may confound the mapping process. We opted to exclude from further analysis any haplotig/primary contig pairs that did not map to the same region on the reference.

### Assessment of genome assembly quality

*Comparison with reference genome:* The dot-plot comparison of our assembled genome and GRCh38 was generated by http://assemblytics.com [27] using the output of mapping the primary contigs to the reference genome using the MUMmer program.

*Content of conserved gene:* The lineage-specific sets of Benchmarking Universal Single-Copy Orthologs (BUSCOs)[28] were evaluated on primary contigs using the BUSCOs 3.0.2, BLAST 2.6, Augustus 3.3, EMBOSS 6.6.0, and HMMER 3.1b2.

*Comparison with other sequencing technology:* Illumina data for HG002 at 71x coverage was mapped to the assembled genome using BWA-MEM version 0.7.12[29,30]. The mapped reads were assessed for mismatch percentage with the Qualimap program version 2.0[31,32].

### Genotyping and identification of informative SNPs from trio short read data

A set of 62x, 70x, and 71x coverage, 250-bp Illumina paired-end data of the father (HG003), mother (HG004), and son (HG002) of the Ashkenazi trio was retrieved from the Genome in a Bottle Consortium[4]. Each pair of FASTQ file was mapped to the GRCh38 human reference genome and genomic variants were called using Sentieon’s variant calling pipeline[33]. Joint genotyping of the three individuals was performed and only the loci that were present in all three individuals and concordant with Mendelian inheritance were selected for further analysis.

An informative SNP was defined as a SNP in the child for which we can unambiguously determine the parental origin. In order to determine parent of origin, we selected heterozygous loci in the child where the parents have different genotypes. For example, the loci with genotype A/T, G/T and A/T in the father, mother, and child respectively, would satisfy this criteria. The A allele in the child must be the paternal SNP and the allele T in the child must be the maternal SNP. As another example consider a genotype of C/C, T/T, and C/T in the father, mother, and child respectively. In this case, the C allele must be the paternal allele while the T allele must be the maternal allele.

### Screening and validating of phasing

Once our set of informative SNPs was generated, we performed a series of filtering steps to concentrate on those primary and haplotype contigs that we can confidently determine phasing accuracy (Table 1). Our filtering steps were as follows:

1. For each haplotype contig and its corresponding section on the primary contig, we infer the parental origin of SNPs along them using the informative SNPs of our trio data. The contigs with no informative SNPs were removed.
2. There are SNPs that should be heterozygous based upon parental inheritance, but are represented as homozygous in the primary contigs and haplotigs. These reflect either errors in FALCON-Unzip’s segregation of the haplotypes or insufficient coverage in the long read data to detect this allele. These errors were noted and then removed from further analysis because they will cause confusion in determining the parent of origin for the primary or haplotype contigs.
3. The haplotig/primary contig pairs that have less than 10 informative SNPs between them were also removed because they lack sufficient resolution to detect or classify phasing errors.
4. The contigs that mapped to multiple chromosomes were removed from further analysis because they are potentially misassembled regions or segmental duplications, which will obscure the interpretation of phasing accuracy.
5. Moreover, we also removed contigs that are interrupted by other contigs when mapped to the reference genome. Because these regions may involve structural variations of the genome, their inclusion would affect our downstream interpretation of the distribution of informative SNPs along haplotigs (Fig 1C).
6. The contigs that mapped to non-autosomes except pseudo autosomal regions on chromosome X were also removed because presumably these regions should come directly from one parent or the other.

We call the haplotigs/primary contigs remaining after this filtering process our “haplotig analysis set”.

**1.**
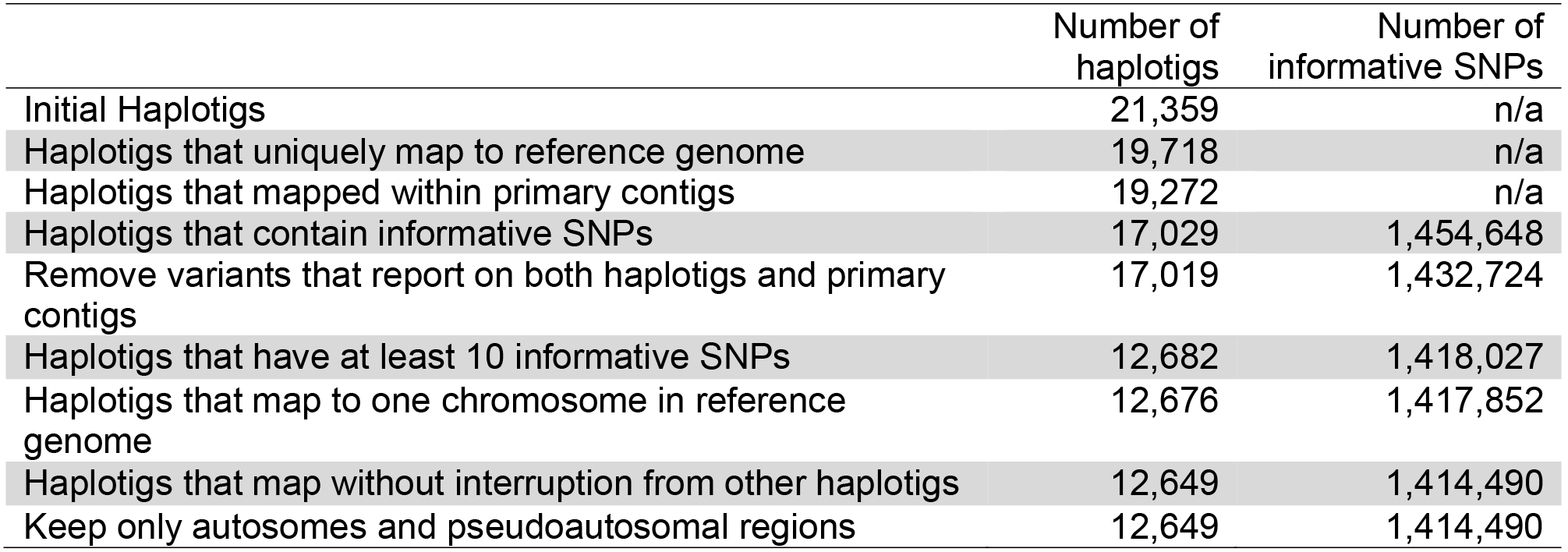
Haplotig filtering statistics

After filtering, we label the parental origin of a haplotig and its corresponding primary contig based on the informative SNPs found along the contigs. The SNPs along those contigs that have a parental origin different from the norm would be considered as mis-phased SNPs. For example, if a haplotig contains 11 informative SNPs from the mother and two informative SNPs from the father and the corresponding section on the primary contig contains seven informative SNPs from the father and one informative SNP from the mother, we will classify the haplotig as the maternal contig and the corresponding section on the primary contig as the paternal contig. Therefore, the two informative SNPs on the haplotig and one informative SNP from the corresponding primary contig are considered as mis-phased SNPs.

Each haplotig and its corresponding section on the primary contig was analyzed for the percentage of mis-phased SNPs, the locations of mis-phased SNPs, the number of continuous mis-phased SNPs, and the minimum number of recombination events that could rearrange all mis-phased SNPs into the correct parents, a metric referred to as the “switch error” [17].

## RESULTS

### Genome assembly quality

The assembled, phased, and polished genome of HG002 consisted of 1,376 primary contigs containing 2.8 Gbp with an N50 contig length of 8.9 Mbp and a maximum contig size of 36.7 Mbp (Table 2). These assembled contigs spanned the majority of the GRCh38 human reference genome except the centromeric regions (Fig 2). To further assess the quality of the genome assembly, we make use of the Benchmarking Universal Single-Copy Orthologs (BUSCOs) gene set. BUSCO genes are highly conserved genes across the given lineage, and tracking the percentage of BUSCO genes present in an assembly can provide additional information about the assembly’s integrity and completeness. Our assembled contigs contained 2,399 complete and 2,368 complete and single-copy BUSCO groups out of 2,586, which corresponds to 92.7% and 91.5% respectively (Table 3). This evidence suggests that our assembled genome covers the majority of the human genome.

**2.**
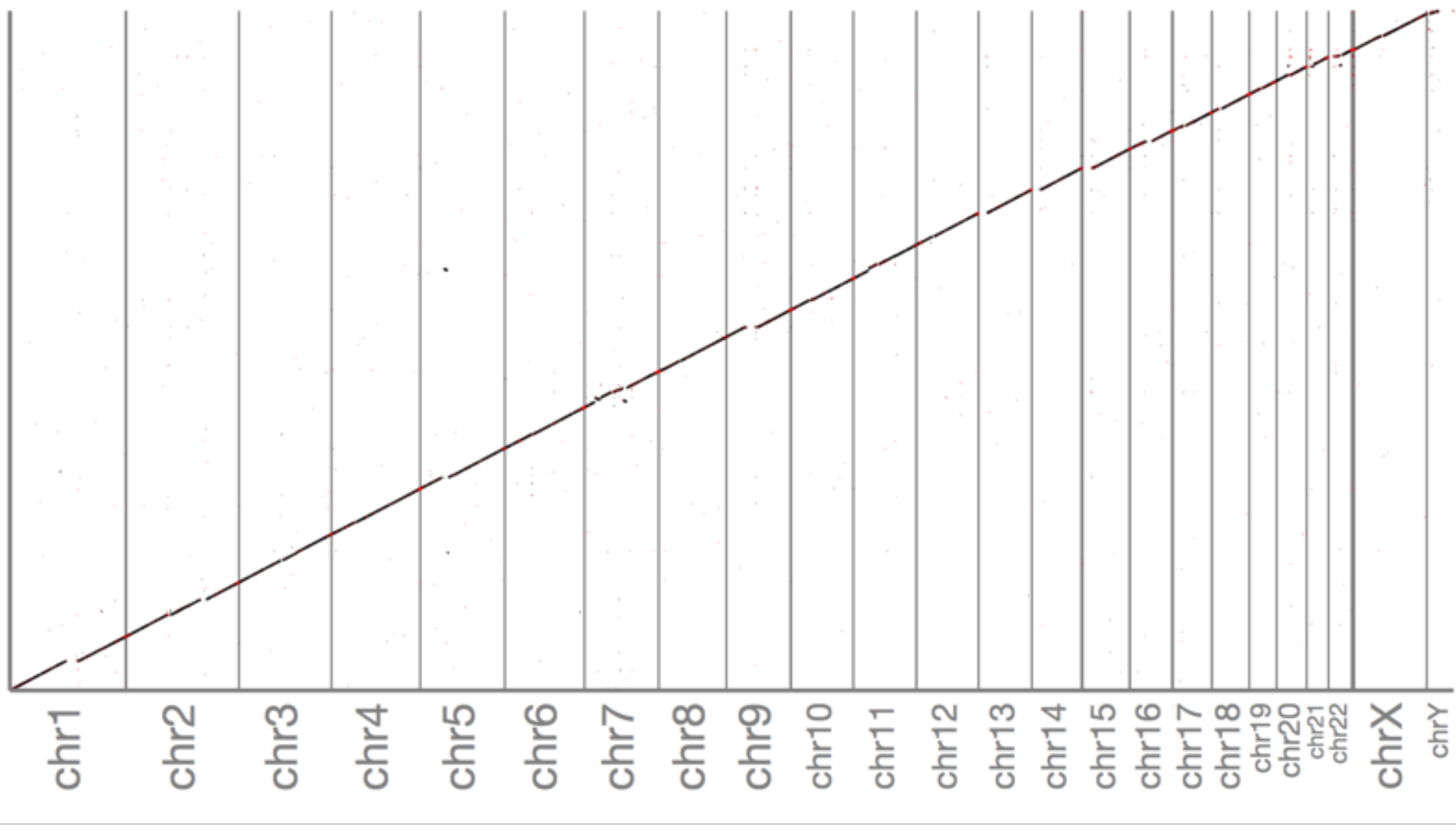
Dot plot of assembled primary contigs against GRCh38 human reference genome.

**2.** Assembly statistics

2 Assembly statistics
|  | Number of<br>bases | Number of<br>contigs | N50 | N90 | Largest<br>contig |
| --- | --- | --- | --- | --- | --- |
| Raw reads | 219,010,844,467 | 26,612,068 | 11,094 | 4,782 | 88,912 |
| Error corrected<br>reads | 112,526,506,703 | 15,109,954 | 10,249 | 4,315 | 47,127 |
| Primary assembled<br>contigs | 2,798,120,340 | 2,707 | 8,925,002 | 1,391,709 | 36,640,311 |
| Polished primary<br>assembled contigs | 2,767,334,372 | 1,376 | 8,923,167 | 1,463,789 | 36,674,135 |

**3.** BUSCOs summary score

3 BUSCOs summary score
|  |  |
| --- | --- |
| Complete BUSCOs | 2,399 |
| Complete and single-copy BUSCOs | 2,368 |
| Complete and duplicated BUSCOs | 31 |
| Fragmented BUSCOs | 98 |
| Missing BUSCOs | 89 |
| Total BUSCO groups searched | 2,586 |

In addition, using Illumina data from the same individual, 99.77% of the Illumina reads can map to our assembled contigs with a general error rate of 1.06%. These mappings and mismatch percentages were similar to those found when aligning the same Illumina data to the GRCh38 reference genome, which had 99.9% mappability and 0.79% general error rate. This suggests that our assembled genome has low error rate for the small indel and mismatch combined.

### Genome phasing statistics

The FALCON-Unzip segregated the assembly into 21,359 haplotigs containing 1.68 Gbp with, an N50 contig length of 136 kbp, a N90 contig length of 34 kbp, and a maximum haplotig size of 1.36 Mbp. Although, there were 3.18 million heterozygous SNPs in the Ashkenazi son, only 2.43 million of them were informative SNPs, and about 1.45 million of them were informative SNPs located inside haplotigs/primary contigs. This number of informative SNPs in haplotigs/primary contigs is expected given that there are 2.43 million informative SNPs in 2.8 Gbp primary contigs. If the informative SNPs are uniformly distributed, the 1.68 Gbp regions that have haplotigs should have 1.68*2.43/2.8 = 1.458 informative SNPs.

These contigs were filtered to retain only haplotigs that we can reliably estimate phasing accuracy using their mapped locations and informative SNPs from Illumina data (See above in Coordinating and calling variants of primary and haplotype contigs and Screening and validating of phasing in the Materials and Method section). The largest proportion of haplotigs were removed due to having less than 10 informative SNPs (Table 1). After filtering as described above, we were left with around sixty percent of haplotype contigs segregated by FALCON-Unzip that have sufficient information to be included in this evaluation. The haplotig analysis set includes 1,414,490 SNPs along 12,649 haplotigs covering 1.2 Gbp. On average, we find informative SNPs every 850 bp on either a haplotig or its corresponding section of the primary contig. The informative SNPs are found in roughly similar numbers on the primary contigs (644,876 SNPs) and haplotigs (769,614 SNPs) or between father (706,053 SNPs) and mother (708,437 SNPs).

### Genome phasing accuracy

Overall, 9,212 of our haplotig analysis set (72.8%) contained no mis-phased SNPs along the haplotigs or their corresponding sections on primary contigs, which suggests that they are accurately phased (Fig 3A). These haplotigs covered 776 Mbp of the reference genome and corresponds to 64.3% of the base pairs in the haplotig analysis set. The longest haplotig spanned up to 1.05 Mbp and contained up to 1,266 SNPs.

**3.**
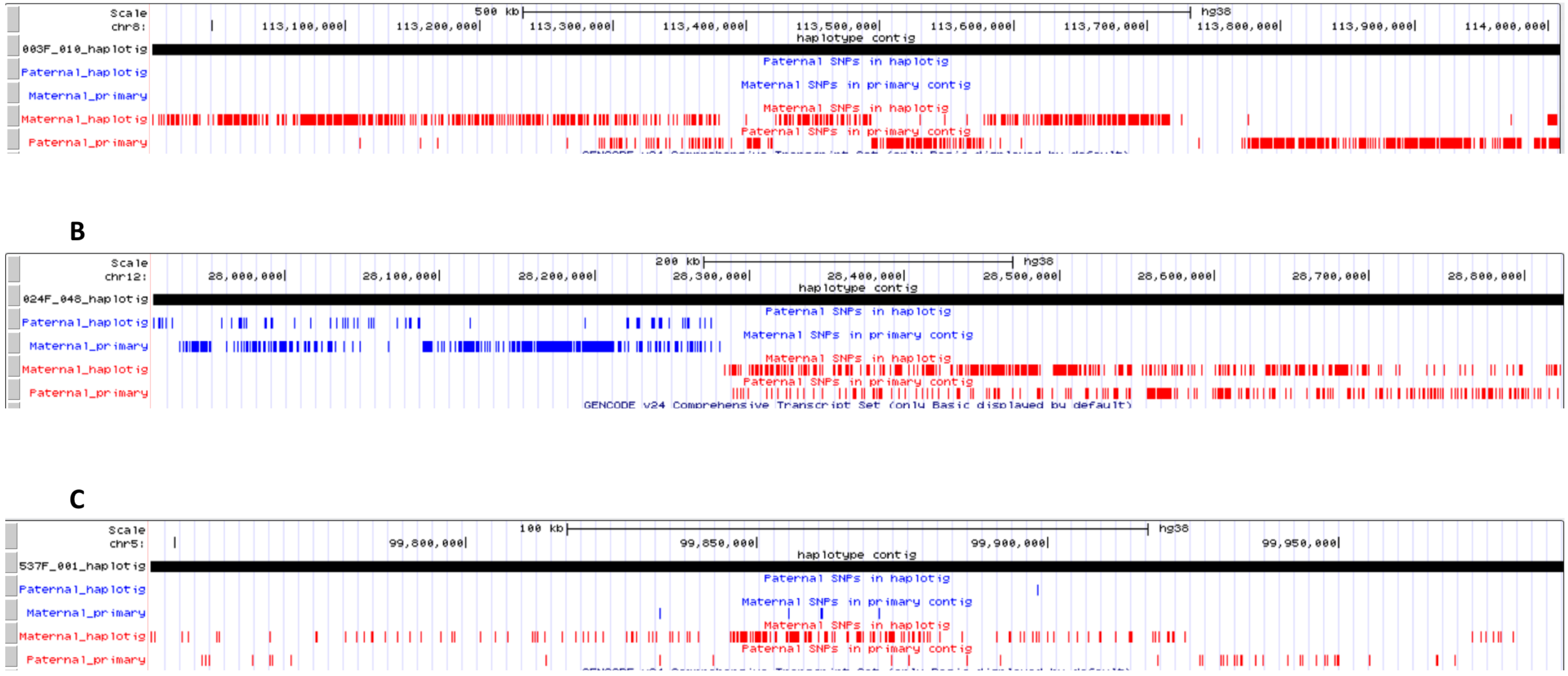
Distribution of informative SNPs along haplotigs and corresponding primary contigs. The screenshots were taken from UCSC genome browser. A) correct phasing haplotig B-C) incorrect phasing haplotigs: mis-joined haplotigs and random error haplotigs

The remaining 3,437 haplotigs in our analysis set (27.2%) contained at least one mis-phased SNP (Fig 3B, 3C). These haplotigs covered 430 Mbp of the reference genome and contained between 0.0007-50% mis-phased SNPs. The longest mis-phased haplotig spanned up to 1.15 Mbp and contains up to 424 SNPs. Among these haplotigs, 2,043 (covering 219 Mbp) contained only a single mis-phased SNP and 2,656 (covering 320 Mbp) had mis-phased SNPs comprising less than 10% of the number of informative SNPs.

### Mis-phased pattern

When examining the patterns of the number of mis-phased SNPs, we observe both highly clustered and sparse mis-phased SNPs. Such patterns are important to be further categorized because the clustered mis-phased SNPs are more likely to be generated from mis-joining two haplotigs, while the sparse mis-phased SNPs are more likely to be generated by assembly errors, local haplotype assignment errors, or genotyping errors. In cases where we see five or more consecutive informative SNPs coming from the other parent, we designate these errors as “mis-joined errors”. In cases where we see two or fewer consecutive informative SNPs coming from the other parent, we designate these errors as “random error errors.” In all of the 3,437 haplotigs with non-perfect parental segregations, we found 588 haplotigs contain only misjoined errors which are labeled “mis-joined haplotigs” while 2,506 haplotigs contain only random errors which are labeled “random error haplotigs” (Fig 4). As for the rest, 125 contigs contain both random and mis-joined errors in the same contigs and 218 contigs contain three to four consecutive informative SNPs coming from the other parent. In this research, we explore the characteristic of mis-joined haplotigs and random error haplotigs in detail.

**4.**
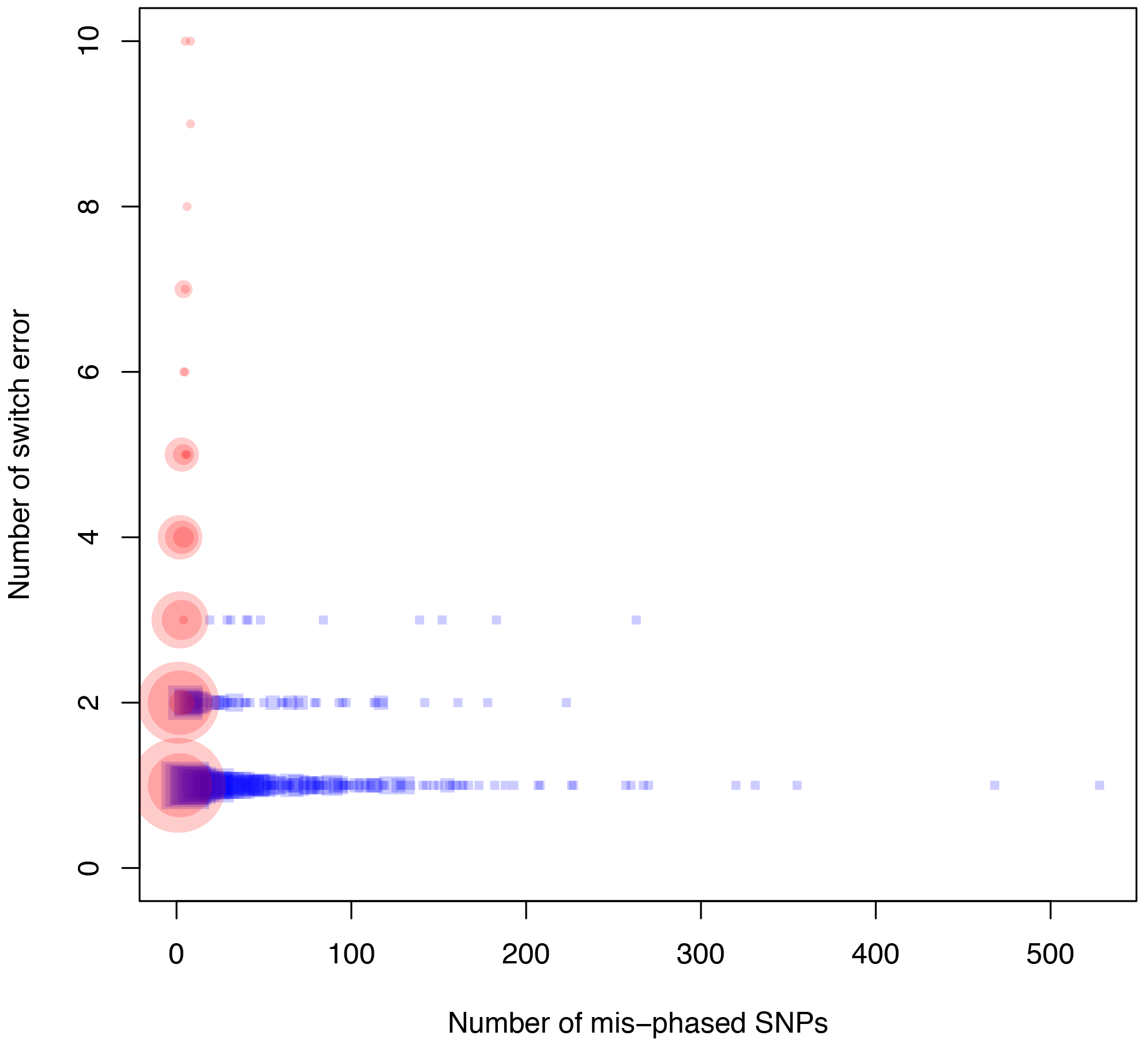
Scatter plot of number of mis-phased SNPs and number of switch error for haplotigs. Blue boxes are haplotigs with minimum cluster size of five mis-phased SNPs (mis-joined haplotigs). Red dots are haplotigs with the maximum cluster size of two mis-phased SNPs (random error haplotigs). The size of the boxes and dots are natural logarithm of number of haplotigs with the same number of mis-phased SNPs and switch errors plus one.

The mis-joined haplotigs (Fig 3B, Blue rectangles in Fig 4) have the following characteristics:

- First, interestingly, the number of switch errors – the minimum number of recombination events that could rearrange all mis-phased variants to the correct parent -- does not appear to increase with the number of informative SNPs (Blue rectangles in Fig 4) which might be expected if each mis-phased SNP offered a random chance at switching parental origin. Instead, this seems to indicate that these errors are typically due to a small number of mis-joined haplotigs, with long stretches of informative SNPs from the same parent.
- Second, we defined the area between the set of SNPs from one parent and the transition to the set of SNPs from the other parent as the “boundary region”. For misjoined haplotigs, these boundaries are not located specifically in any part of haplotig suggesting that the mis-joined event could occur anywhere along the haplotigs (Fig 5).
- Third, the distances at the boundaries are large with a median of 9,089 bp and interquartile range of 7,440 bp (Fig 6A) as compared to average distance between informative SNPs at 850 bp. These large distances at the error boundaries imply that most mis-joined events occurred in low SNPs density regions.

**5.**
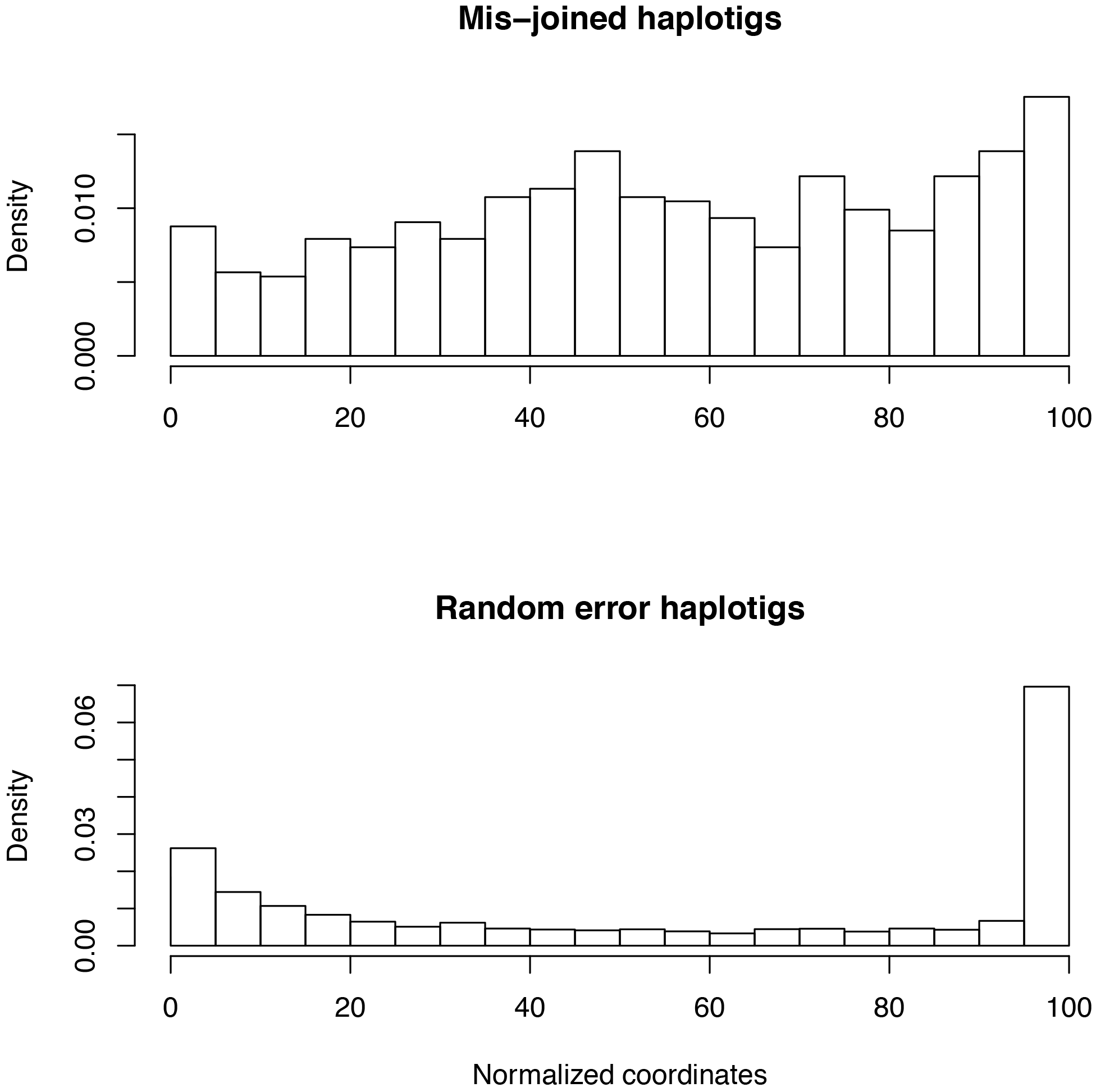
Histograms of normalized locations of mis-phased boundary. The locations of boundaries are assigned to the right SNPs which lead to right skew of the histogram. The random error haplotigs (lower figure) have high density of mis-phased boundary toward ends of haplotigs

**6.**
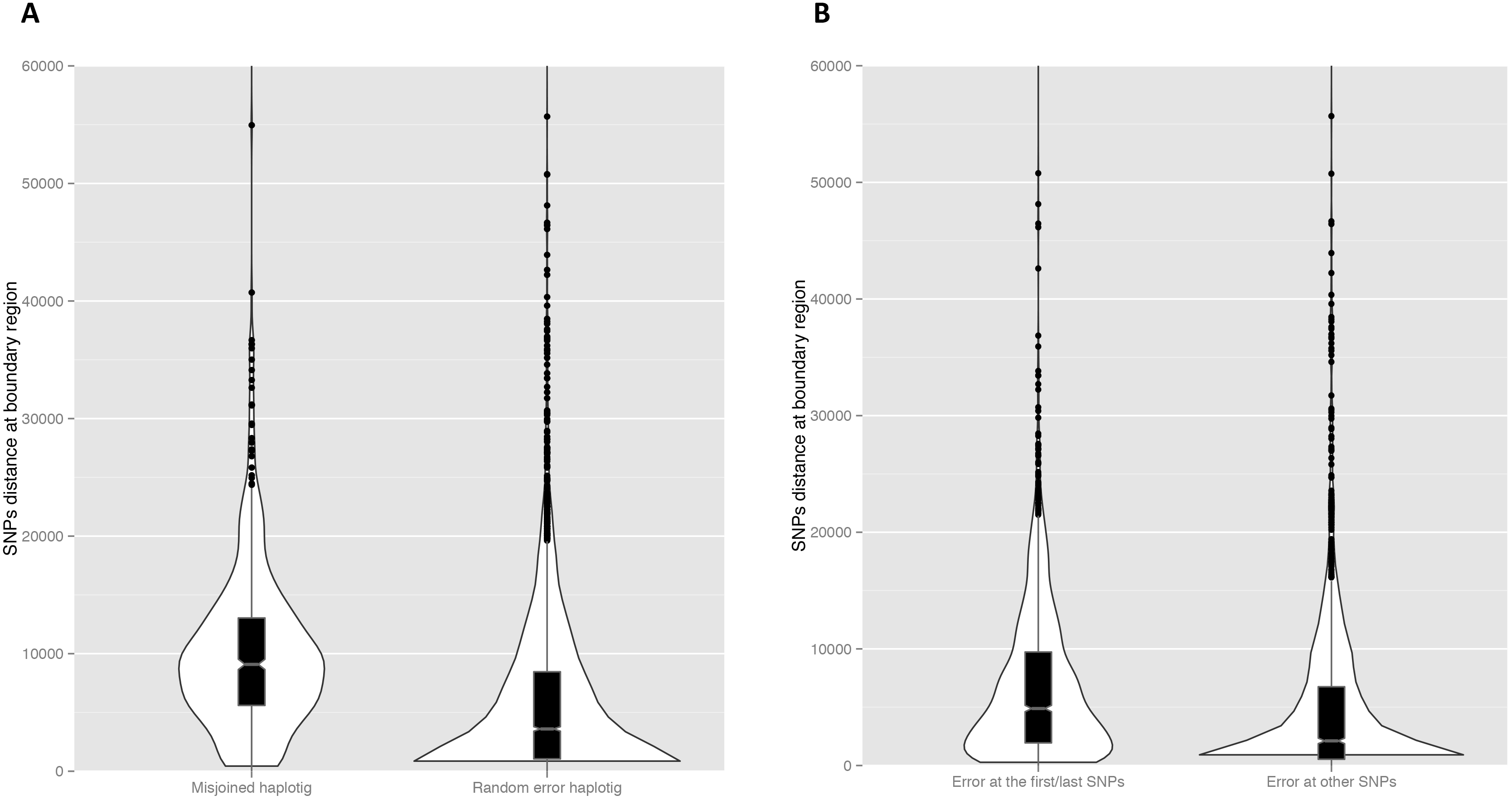
Box plots and violin plots of distance at boundary between correct and mis-phased SNPs. A) comparison between mis-joined and random error haplotigs B) comparison between errors at the first/last informative SNPs of the contigs and the SNPs in other locations for random error haplotigs

In contrast, the random error haplotigs (Fig 3C, Red dots in Fig 4) have the following characteristics:

- First, the number of switch errors linearly increases with the number of mis-phased SNPs. This characteristic is due to the fact that instances with more consecutive mis-phased SNPs were classified as mis-joined haplotigs.
- Second, the boundaries between correct and mis-phased SNP cluster toward the end of haplotigs (Fig 5).
- Third, the distances of boundaries between correct and mis-phased SNPs in random error haplotigs is significantly shorter (*P*<2.2e-16: Mann-Whitney-Wilcoxon test) than those of mis-joined haplotigs. Most of the boundaries are short (violin plot Fig 6A) with a median of 3,598 bp and interquartile range of 7,412 bp. Due to the prevalence of random error SNPs, we observe such errors in both high and low SNP density regions. Interestingly, the detailed analysis of random error haplotigs shows that such distances of boundaries that involve the first/last informative SNPs are longer than those that do not involve the first/last informative SNPs (*P*<2.2e-16: Mann-Whitney-Wilcoxon test) (Fig 6B).

## DISCUSSION

### Quality of genome assembly using FALCON

The FALCON genome assembly of the Ashkenazi son captures a significant proportion of the true human genome based on a comparison to the most recent human reference genome and a list of evolutionarily informed expected genes.

The N50 contig size in this study (8.9 Mbp) is longer than the first diploid human genome assembly using shotgun sequencing (106 kbp)[34] and the first long-read haploid human genome assembly (144 kbp)[6]. A more recent study was able to achieve a contig N50 of 17.9 Mbp in a diploid human sample using a combination of long reads, short reads, and microfluidics-based linked reads[35]. Moreover, a recent assembly produced using Oxford Nanopore data was able to achieve an N50 contig length of 24.5 Mbp. As technology continues to improve, we anticipate that we will continue to achieve higher continuity assemblies with quality comparable to the well-defined reference genome[13].

### Quality of genome phasing FALCON-Unzip

The FALCON-Unzip is capable of segregating the assembled genome into long haplotigs with a small proportion of haplotigs that were mis-joined. Compared to the original FALCON-Unzip publication which looked at a Arabidopsis assembly [3], our human haplotigs are shorter (N50=136 kbp vs 6.9 Mbp), but a smaller proportion of them contain mis-phased SNPs (28% vs 65%) than those of Arabidopsis (Table S5 from Chin 2016 publication[3]). The difference in ratios of mis-phased contigs could come from the stringency of filtering methods or the different genome architectures and data between two species.

We observe two types of mis-phased haplotigs: the mis-joined haplotigs and the random error haplotigs. Interestingly, there is less evidence of mis-haplotig in the original FALCON-Unzip publication which verifies the algorithm using a cross of two inbred Arabidopsis specimens. The Arabidopsis genome is simpler than the human genome in terms of the number of chromosomes, genome size [36], and percentage of repeat content[36,37]. Such factors could make haplotype assembly for humans more challenging and lead to the higher abundance of mis-joined haplotigs reported in this publication.

The observation that most mis-phased SNPs in random error haplotigs are clustered toward the end of haplotigs (Fig 5) is intriguing. One possible explanation is that that the algorithm, which determines the boundary of a haplotig, has made inexact cuts and consequently the haplotig includes some component of other haplotypes. One observation supporting this hypothesis is that when we encounter a misphased SNP as the first or last SNP of a haplotig, it is much more likely to be found in an area of low SNP density (Fig 6B). The effect of incorrect allele assignment around low SNPs intensity regions could affect both the decision to join haplotigs and the accuracy of haplotig boundaries. We suggest that FALCON-Unzip should explore increasing the stringency of parameters used in terminating a phase block.

## CONCLUSION

The advancement of genomic technologies and availability of massive computing resources have transformed our paradigm of scientific procedure. In this publication, we demonstrate that we can both assemble AND phase the human genome with high quality in a single experiment, while it took a global effort a decade to unveil an unphased, more fragmented first draft of the human genome in 2001. FALCON-Unzip can phase 72 percent of haplotigs with no error and the majority of errors are random. The low SNPs density area would need to be inspected as they are the hotspot of mis-joined error. As the algorithm continue to improve, we predict that this framework could achieve higher overall accuracy and lead to many more high quality de novo genome assembled and phased genomes.

The applets for data analysis in this research is publicly available on DNAnexus and GitHub https://github.com/Arkarachai/falconunzipassessment. These applets generated all the results, statistical analyses, and figures in this manuscript.

## ACKNOWLEDGEMENTS

We would like to thank Nicholas Hill, Eric Talevich, and Samantha Zarate for their comments on the manuscript. This project is fully supported by DNAnexus for computing and storage. The DNAnexus company disclaims responsibility for any analyses, interpretations, or conclusions.

